# Assembly of Mb-size genome segments from linked read sequencing of CRISPR DNA targets

**DOI:** 10.1101/373142

**Authors:** GiWon Shin, Stephanie U. Greer, Li C. Xia, HoJoon Lee, Jun Zhou, T. Christian Boles, Hanlee P. Ji

## Abstract

We developed a targeted sequencing method for intact high molecular weight (**HMW**) DNA targets as large as 0.2 Mb. This process uses HMW DNA isolated from intact cells, custom designed Cas9-guide RNA complexes to generate 0.1 – 0.2 Mb DNA targets, electrophoretic isolation of the DNA targets and sequencing with barcode linked reads. We used alignment methods as well as local assembly of the target regions to identify haplotypes and structural variants (**SVs**) across multi-Megabase genomic regions. To demonstrate the performance of this approach, we designed three assays that covered a 0.2 Mb region surrounding the *BRCA1* gene, a set of 40 overlapping 0.2 Mb targets covering the entire 4-Mb MHC locus, and 18 well-characterized structural variants. Using the highly characterized NA12878 genome, we achieved on-target coverage of more than 50X, while overall whole genome coverage was approximately 4X. We generated haplotypes that completely covered each targeted locus, with a maximum size of 4 Mb (for the MHC region). This method detected structural variants such as deletions and inversions with determination of the exact breakpoints and genotypes. Even breakpoints inside highly homologous segmental duplications are precisely determined with our high-quality assemblies. Overall, this is a new method to sequence large DNA segments.

---

Sequencing intact DNA molecules of high molecular weight (**HMW**) provides insight into complex genomic features such as haplotypes and structural variants (**SVs**) that span many kilobases (**kb**) to Megabases (**Mb**). Frequently, these features are obscured when one uses short reads that come from libraries where DNA is fragmented to insert sizes less than 1kb. To leverage the advantages of long-range sequencing of HMW DNA molecules, some instruments provide long reads such as Pacific Biosciences (Menlo Park, CA)^1^ and Oxford Nanopore Technologies (Oxford, UK)^2^, or by linked read sequencing using 10X Genomics (Pleasanton, CA) library construction^3–5^. These approaches have been used in whole genome sequencing (**WGS**). However, WGS methods have practical limits based on the extent of sequencing coverage, higher cost per sample and the complexities of analyzing large data sets representing the full genome.

There is increasing interest in targeted sequencing of HMW DNA as an alternative to WGS. High coverage of intact HMW targets may improve the characterization of complex genomic features such as candidate structural variants (**SVs**), contiguous haplotypes and assemblies for regions of interest^1,2,5,6^. The benefits of higher targeted coverage are particularly relevant in the context of genetic mixtures where SVs or other variants are present in a low allelic fraction. In addition, sequencing of target HMW DNA molecules offers a means of evaluating regions-of-interest more efficiently and cost-effectively than with WGS. This targeted approach has potential implications for diagnostic tests currently reliant on low-resolution methods such as fluorescent in-situ hybridization (**FISH**).

We describe a new approach for targeted sequencing of HMW DNA molecules (**Fig. 1a**). This method has several discrete steps. First, we isolate intact chromosome-length HMW DNA from intact cells using a rapid gel electrophoresis method. The extracted HMW DNA is then digested with specific Cas9-guideRNA complexes that excise the genomic sequencing targets (up to 0.4 Mb). Referred to as Cas9-Assisted Targeting of Chromosome Segments (**CATCH**)^7^, the target DNA is substantially enriched from off-target molecules with size-selection electrophoresis – this step eliminates a significant fraction of off-target molecules in different size ranges (**Supplementary Fig. 1**). The extraction, Cas9 digestion, and size selection are automated on a SageHLS instrument (Sage Science). Guide RNAs (**gRNAs**) are synthesized with a conventional phosphoramidite method, or using highly multiplexed *in vitro* transcription reactions with DNA templates produced by oligonucleotide array technologies.

**Figure 1.**
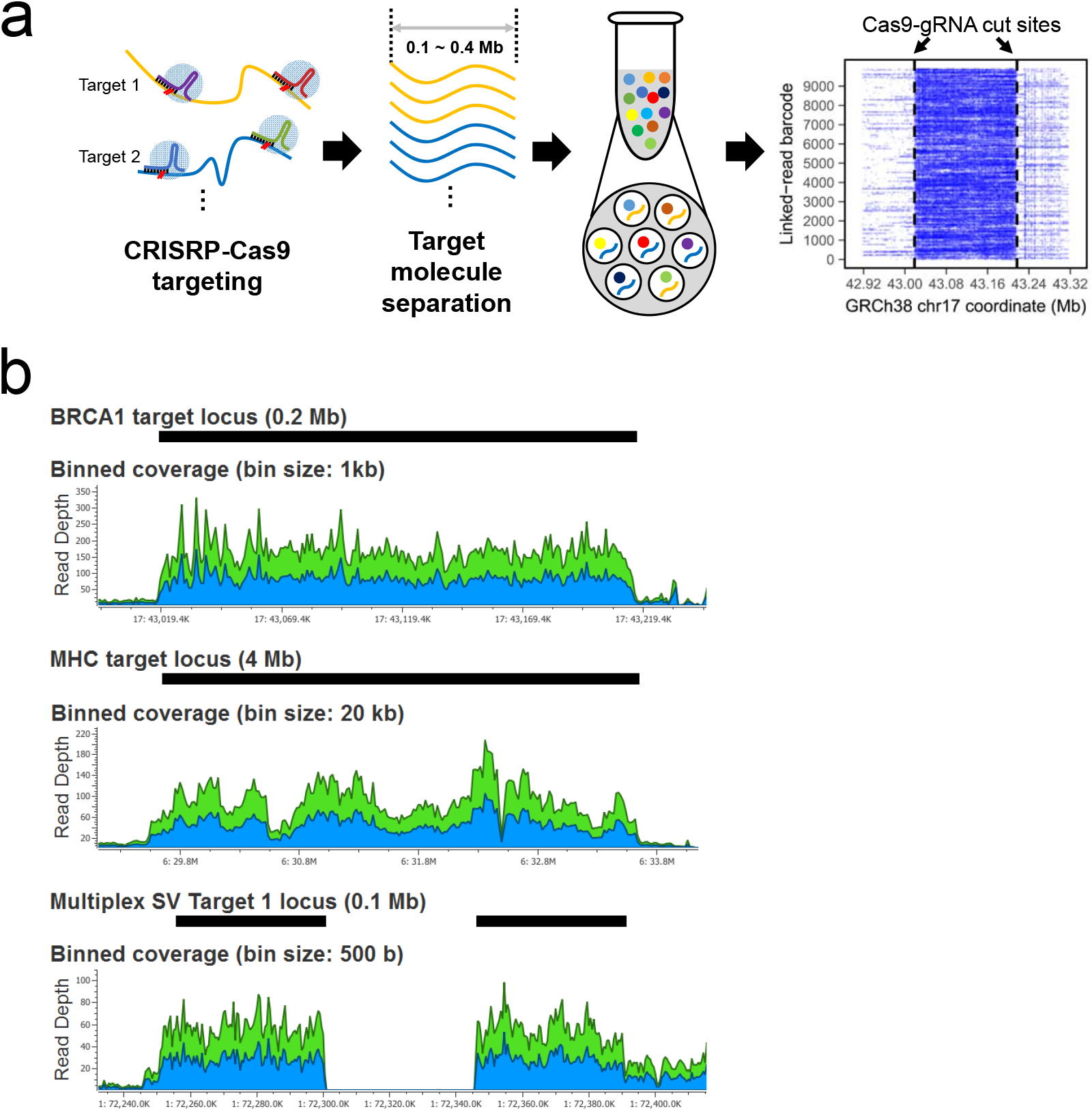
CATCH targeting and linked read sequencing of HMW DNA. **(a)** Overview of the process is illustrated. First, guide RNAs target and cut multiple genomic regions of interest. Second, target HMW DNA within the specific size range is isolated by an electrophoresis-based process. Lastly, the target DNA is used for linked read library preparation and sequencing. The alignment of barcode linked reads shows how sequence coverage is increased across the target segment. In the alignment plot, the X axis indicates the reference coordinates and the Y axis shows different barcodes representing individual HMW molecules. Dashed vertical lines indicate Cas9-gRNA cut sites. **(b)** Sequencing coverage for the target regions are shown for the three assays. For Assays 1 and 2, *BRCA1-R2* and MHC-30 libraries are shown. For Assay 3, an example of a homozygous deletion is shown. Black bars indicate the target regions. Blue and green areas in plots indicate coverages for forward and reverse reads respectively.

Second, the HMW DNA targets are processed into a linked read library with an automated Chromium microfluidics platform (10X Genomics). The HMW DNA is distributed across approximately one million droplets, each with an oligonucleotide barcode reagent. Only a small number of molecules, typically three to five, are present in each droplet. A linear amplification with the HMW DNA template occurs in each droplet. This step produces a synthetic DNA molecule that incorporates the droplet-specific barcodes. After sequencing, one uses the barcodes to link specific reads to individual HMW DNA molecules. In our previous studies, we have used linked read sequencing to define highly complex structural variations and rearrangements across Mb-length genome regions that otherwise may have been missed with conventional sequencing methods that rely on short DNA inserts^3–6^.

Third, we use an Illumina sequencer to generate high sequencing coverage from the linked read targeted libraries. The barcodes link individual reads to their originating HMW DNA molecules, present within a specific droplet. This bioinformatics process involves an alignment-based approach to delineate extended variants, haplotypes and SVs as well as assembly methods for producing scaffold from target regions (**Supplementary Fig. 2**).

For this study, we designed three assays and evaluated their performance on DNA from NA12878, a well-characterized genome that has been sequenced across multiple platforms. We assessed the accuracy of this method to characterize haplotypes and structural variants (**SVs**) across Mb-size targets. Guide RNAs were designed to cut 0.1 to 0.2 Mb targets, either individually or in a tiled fashion. The assays targeted the following: i) a 0.2 Mb region encompassing the entire *BRCA1* gene locus; ii) a 4 Mb segment containing the entire Major Histone Compatibility (**MHC**) locus; iii) 18 SVs of different classes present within 0.1 Mb segments (**Supplementary Table 1**). After DNA preparation, we conducted linked read sequencing and data analysis.

For the first assay targeting the entire *BRCA1* locus on the long arm of chromosome 17 (17q21.31), we ran two replicate experiments (*BRCA1-R1* and *BRCA1-R2*) with NA12878. Both replicates provided adequate yields of target DNA for linked read library preparation without any intermediate processing steps. We observed a 16-fold and 36-fold increase for sequences from the intact *BRCA1* DNA segment compared to off-target sequences across the two assays (**Fig. 1b, Supplementary Table 2**). The fold increase of on-target *BRCA1* segment DNA was concordant with qPCR copy number measurements (**Supplementary Table 2, Supplementary Fig. 3**). Moreover, the parental haplotypes obtained by the alignment-based approach (**Online Method**) were 100% concordant with the previously described NA12878 phased variants (**Supplementary Table 3**).

We used the Supernova program^8^ to generate assemblies with the linked read sequences from the target *BRCA1* region and all other assays. Each assembly used the on-target linked reads (**Online Methods**). For the *BRCA1-R1* replicate, the assembly-generated scaffold was 0.2 Mb, nearly the same size as the original target. The scaffold haplotypes were concordant with reported haplotypes^9^ and long read assemblies^1,2^ (**Supplementary Table 4, Fig. 2a**). For the *BRCA1-R2* replicate, the assembly scaffold was 0.148 Mb which was slightly smaller than the original target DNA as a result of gaps (**Supplementary Table 4**).

**Figure 2.**
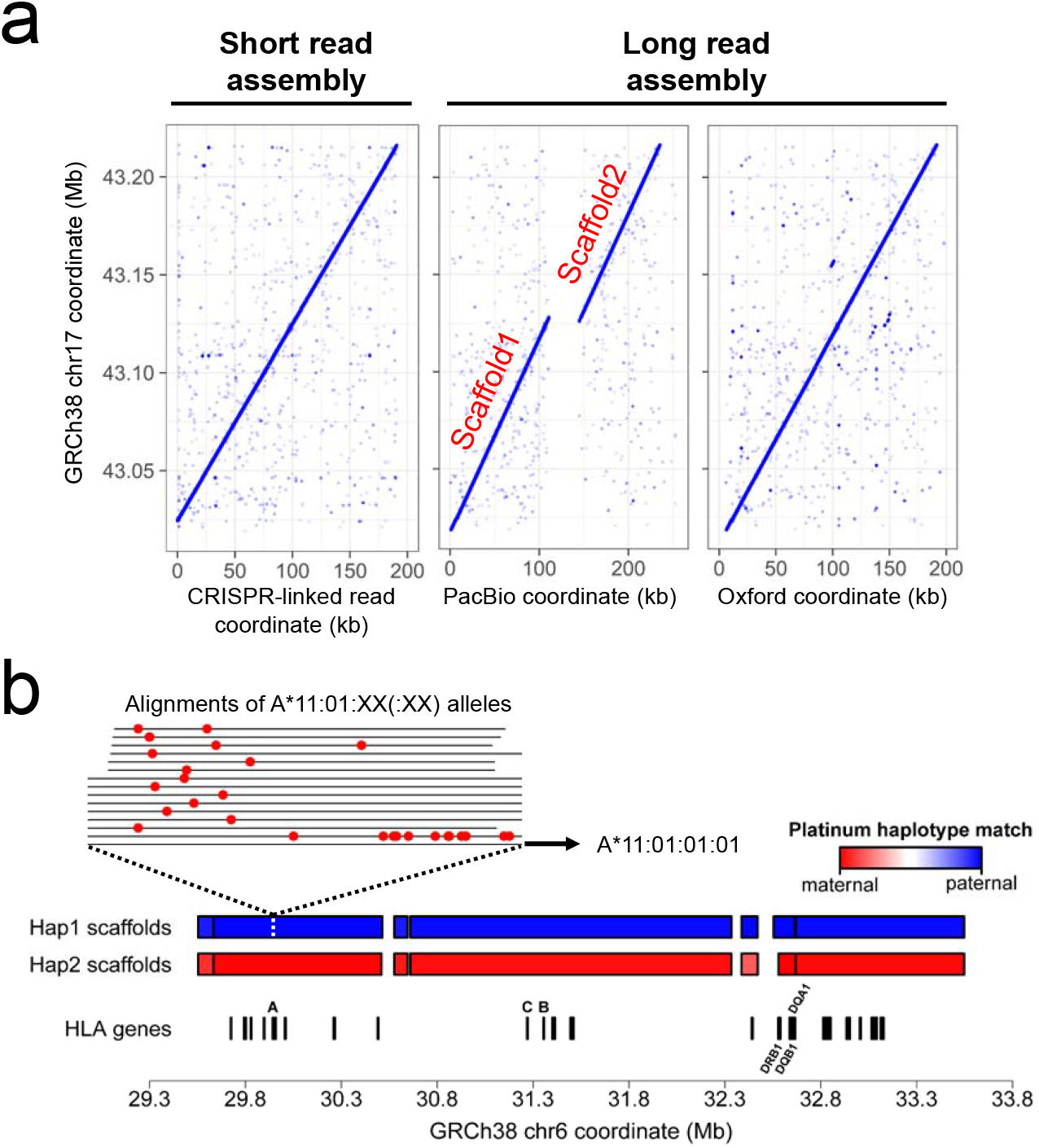
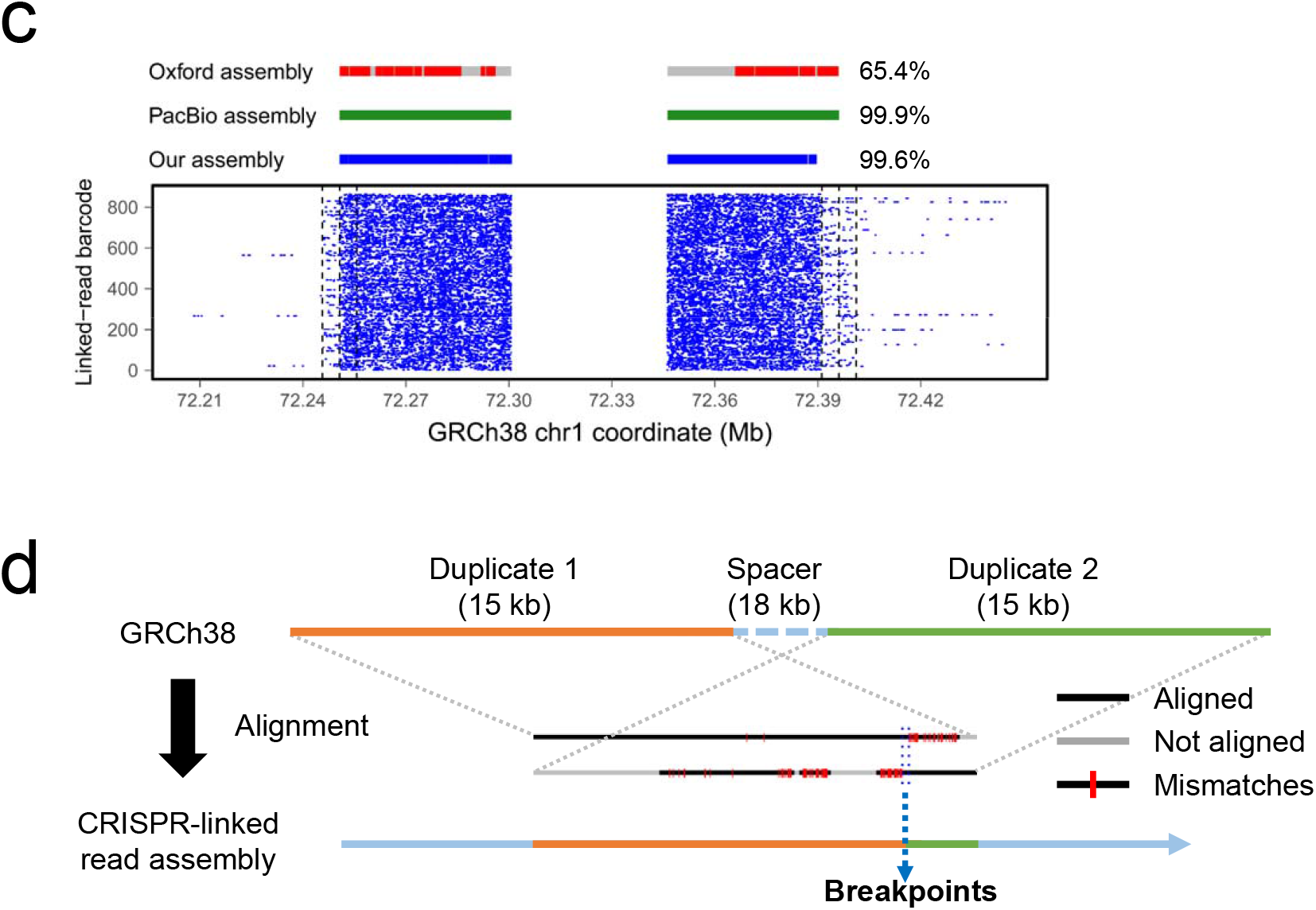
Assembly results. **(a)** The 0.2 Mb *BRCA1* assembly (BRCA1-R1) was compared with other long-read assemblies. The X axis indicates the coordinates of each NA12878 assembly across different platforms. The Y axis indicates the corresponding segment from the GRCh38 reference. Each point indicates where a CRISPR-linked read aligned to the reference versus where it aligned to the NA12878 assemblies. **(b)** The assembled MHC scaffolds with assigned haplotype blocks where red and blue indicate the parental haplotype. The X axis represents the GRCh38 reference genome on which the assembly scaffold is aligned. The HLA genes are indicated below the scaffolds, with the labels only for the major class I (*HLA-A, HLA-B*, and *HLA-C*) and II (*HLA-DRB1, HLA-DQA1*, and *HLA-DQB1*) genes. For *HLA-A*, alignment of all the alleles coding the same protein sequence [A*11:01:XX(:XX)] is shown. The red dots indicate mismatch bases to the allele in Haplotype 1 (A*11:01:01:01). **(c)** CRISPR-linked read assembly for SV1 in Assay 3 was aligned to the reference genome and compared with other long read assemblies. The X axis indicates the reference coordinates. Red, green and blue bars indicate the portion of the assembly that aligns to the reference while a grey gap indicates no alignment. Although the grey regions have some similarity to the reference, they generally have too many homopolymer errors resulting in alignment failure. Fraction of aligned bases in the assembly is indicated at the end of the bars. In the bottom panel, the Y axis shows the alignment of barcode linked reads from SV1. Dashed vertical lines indicate Cas9-gRNA cut sites. **(d)** Illustration of how the breakpoints are determined in segmental duplications. Duplicated copies from GRCh38 reference genome were aligned to CRISPR-linked read assemblies. According to their alignment and mismatches, breakpoint ranges in the reference duplicates were determined. The example shown here is from our SV17 assembly.

Given the difference in scaffold size between the two replicates, we evaluated two methods of downsampling to improve the assembly quality and size: i) random downsampling of a subset of on target linked reads and ii) elimination of barcodes with a higher likelihood of having overlapping DNA molecules with different haplotypes in the same droplet (**Online Methods**). Using the total number of assembled bases excluding gaps (i.e. N bases) as a criterion, both methods improved the net assembly size (**Supplementary Fig. 4a**). For both downsampling approaches, the optimal on-target coverage range was 60-80X, which produced both a high assembly contiguity and phasing quality (**Supplementary Table 4, Supplementary Fig. 4b**). Subsequently, we used 70X coverage to perform assembly on other targets.

For the second assay, our goal was to characterize haplotypes larger than the 0.2 Mb DNA molecule size. We focused on the entire MHC locus spanning 4 Mb on the short arm of chromosome 6 (6p21.3) (**Supplementary Tables 1 and 2**). MHC is a highly polymorphic region of the genome^10^. We designed and used two sets of pooled gRNAs, referred to as Sets 1 and 2 (**Supplementary Fig. 5a**). Both sets generated tandem tiled cuts in 0.2 Mb intervals across the 4 Mb MHC locus. The two gRNA sets were designed with an offset, such that the Set 1 cut sites were separated by 0.1 Mb from the Set 2 cut sites. This strategy generated overlapping segments that were critical for phasing and assembly. Two separate CATCH processes were performed, i.e. one for each pooled gRNA set. The CRISPR-DNA fragments were pooled to prepare a linked-read sequencing library.

To optimize phasing and assembly, we tested different read per barcode ratios for linked-read sequencing. From the prepared linked read sequencing droplets, we made a 90-μl aliquot (MHC-90) with more droplets and a 30-μl volume portion with fewer droplets (MHC-30) (**Supplementary Table 2**). Loaded with a same input amount for sequencing, MHC-90 had twice as many barcodes compared to MHC-30. Both libraries provided 80X on-target average coverage (**Fig. 1b**). Therefore, MHC-30 had a higher read coverage per barcode ratio than MHC-90. Thus, one can extrapolate that MHC-30 had more reads per molecule.

With the alignment-based approach (**Online Methods**), both MHC-90 and MHC-30 provided high quality haplotypes throughout the entire 4-Mb region with a high correlation to the Platinum Genome haplotypes. Overall, we observed a 96% sensitivity and greater than 98% concordance (**Supplementary Table 3**). Moreover, in the assemblies generated by both the libraries, the N50 scaffold sizes were consistently greater than 0.88 Mb. However, fewer barcodes (MHC30) provided a higher quality assembly in terms of contiguity of the scaffold (**Supplementary Fig. 6**).

We assigned haplotypes to the assemblies (**Online Methods; Supplementary Fig. 2**). We identified genotypes for over 27 HLA genes (**Supplementary Table 5**). The HLA genotypes were phased into haplotypes encompassing the entire 4 Mb region (**Fig. 2b**). Particularly, for the six major MHC class I and II genes, MHC-30 generated complete parental haplotypes. Interestingly, the results for the MHC-90 *HLA-DQB1* gene lacked one of the parental haplotypes (**Supplementary Table 5**). Our conclusion from this result was that a limited number of barcodes (as in MHC-30) provided higher quality assemblies. In addition, for the major MHC genes, predictions for parental haplotypes are available from Oxford assembly^2^. The predictions, however, provided a resolution that distinguished variations only in the coding regions. Our assembly-based haplotypes obtained from MHC-30 library matched the predictions, and additionally provided the intronic variations that NA12878 may have.

The third assay was used for multiplexed characterization of SVs previously reported in NA12878 (**Supplementary Table 6**). We selected a ground truth set of 18 SVs previously reported using two or more WGS approaches – either with linked reads^5^ or long reads with Pacific Biosciences^1^ or Oxford Nanopore^2^. These SVs had one or more reported assemblies that covered their location. We designed a multiplexed set of gRNAs that cut sites flanking each SV, encompassed within a 0.1 Mb segment. The SVs included 13 deletions in varying size from 30kb to 150kb, including one homozygous deletion (**Fig. 1b**), and five inversions (**Supplementary Fig. 5b**).

We compared our targeted local assemblies to the reported WGS assemblies based on Oxford Nanopore and Pacific Bioscience sequencing^1,2^. We demonstrated concordant assemblies for 14 of 18 SVs (77.8%) while Pacific Biosciences had 13 (72.2%) and Oxford Nanopore had 15 (83.3%). Our CRISPR-linked read method and Pacific Biosciences generated assemblies that aligned to the reference genome with more coverage than the Oxford assembly (**Fig. 2c**). One inversion event on Chromosome 12 (SV15) did not generate an assembly scaffold due to inefficient Cas9-gRNA cleavage activity. Nevertheless, alignment of the barcode-specific reads of the locus showed both breakpoints of the inversion (**Supplementary Fig. 7**).

Most of the breakpoints identified by our CRISPR-linked read assemblies matched breakpoints from reported long-read assemblies, but there were some discrepancies (**Supplementary Fig. 8**). SV8 and SV9 had breakpoints that matched the Oxford assembly but not the Pacific Biosciences assembly (**Supplementary Table 7**). For SV9, the Pacific Biosciences assembly had a 2.14kb gap of Ns between the two breakpoints; the N-gap matched our predicted breakpoints. Some minor breakpoint-associated events occurred such as insertions and deletions of several bases (**Supplementary Table 7**). These small indels were not identified from alignment-based analysis and are likely to be the result of microhomologies or non-homologous end-joining^11, 12^. For example, SV6 was reported to have a 7-bp sequence at both breakpoints based on alignment but our assembly reported this sequence at only one breakpoint, suggestive of a microhomology-mediated end-joining^12^. For SV5, we assembled two alleles with different deletion sizes, only one of which was confirmed by the Oxford assembly (**Supplementary Fig. 9**).

Seven SV breakpoint targets had one or more segmental duplications present in the targeted breakpoint segment, of which our CRISPR-linked read analysis was able to generate assemblies for five (**Supplementary Table 8**). To examine these sequences in more detail, we aligned our targeted assemblies to the GRCh38 reference – this genome build is more accurate in annotating genomic duplications than the previous version^13^. We identified the breakpoints for three of the cases: SV7, SV16, and SV17 (**Fig. 2d** and **Supplementary Fig. 10**). For example, SV17 locus in the GRCh reference has a duplication of which two 15-kb segments are 93% similar and separated by an 18-kb spacer. A 15-kb region in our NA12878 assembly had alignments from both the reference segments, by which the breakpoints were able to be determined (**Fig. 2d**). We did not identify breakpoints for two of the five cases (SV12 and SV14) because their duplications had a sequence similarity greater than 99%, thus we assumed that our assemblies had only one of the two identical copies.

In summary, our study successfully demonstrated a multiplex targeted sequencing method that enables targeted assembly of genomic regions larger than 0.1 Mb. This approach benefits from the higher sequence quality of short reads that are linked to target HWM DNA molecules. Our study shows comparable assembly contiguity and better haplotype quality based on single nucleotide variants. With linked reads from multi-Mb target segments we successfully generated assemblies and accurate haplotypes even for highly polymorphic segments of the genome such as the MHC locus. This approach was used to delineate multiple types of SVs with base-pair breakpoint resolution - conventional short read sequencing would likely not delineate these SV structures. For a highly polymorphic genes clustered in a large genomic region, we genotyped a CRISPR-linked read assembly at a resolution that distinguished intronic variations. For future studies, we will evaluate if this approach has utility for detecting clinically actionable rearrangements that are causative for a variety of genetic disorders.

## METHODS

See Online Methods section.

### Accession codes

Sequence data is available at the NIH’s Short Read Archive (SRA) with accession code SRP148930.

## ACKNOWLEDGEMENT

The work is supported by National Institutes of Health [2R01HG006137-04 to HPJ and P01HG00205ESH to GS and HPJ]. Additional support to HPJ came from the Doris Duke Clinical Foundation Clinical Scientist Development Award, Research Scholar Grant, RSG-13-297-01-TBG from the American Cancer Society and a Howard Hughes Medical Institute Early Career Grant. We kindly acknowledge Dr. Anuja Sathe for assistance with tissue culture.

## AUTHOR CONTRIBUTIONS

G.S., H.L., C.B. and H.P.J. designed the experiments. G.S. conducted the experiments (with assistance from J.Z. and C.B. in the BRCA1 study). G.S., C.S. and optimized the isolation process. G.S. developed the analysis algorithms. G.S., L.X. and S.U.G. analyzed the data. G.S., S.U.G. and H.P.J. wrote the manuscript.

## ONLINE METHODS

### Guide RNA design

To design our 20-bp guide RNAs (gRNAs), we considered all 20-bp sequences (20-mers) in the region of our target cut sites. The 20-mers had to occur directly adjacent to a Cas9 binding motif to be included as a candidate gRNA. We compared our candidate gRNA 20-mers to all 20-mers that exist in the reference human genome, and retained only those candidate gRNAs that: i) appeared only once in the human genome, and ii) had few similar 20-mer matches, with either one or two mismatches. Sequence uniqueness was examined with respect to both strands of the human genome. To select optimal 20-mer gRNAs, we also took into account the positions of other genetic variants such as single nucleotide variants, insertion-deletions and structural variants, and the location of the 20-mer relative to genomic features such as exons, introns, repetitive sequences, pseudogenes, methylation sites, microsatellite regions, promoters, highly variable sequences such as T cell receptor hypervariability sites, etc. We selected for synthesis the gRNA 20-mers that best satisfied our criteria.

### *In vitro* guide RNA preparation

Array-synthesized oligonucleotide pools were used as templates to prepare gRNA pools for the MHC and multiplex SV assays (**Supplementary Table 9**). Each oligonucleotide/gRNA consisted of four components: an adapter, a T7 promoter, a target-specific region, and a trans-activating CRISPR RNA (tracrRNA) region. For the MHC assay, a total of 126 gRNAs were prepared for the two 100 kb-offset sets of gRNAs (Set 1 and Set 2); gRNAs in Set 1 and Set 2 had distinct (i.e. set-specific) adapters. For the initial amplification, we added forward primers (5’–GAGCTTCGGTTCACGCAATG–3’ and 5’–CAAGCAGAAGACGGCATACGAGAT–3’) that matched to the set-specific adapter sequences and a reverse primer (5’–AAAGCACCGACTCGGTGCCACTTTTTCAAGTTGATAACGGACTAGCCTTATTTTAACTTGCT ATTTCTAGCTCTAAAAC–3’) complementary to the tracrRNA sequence. Primers were chemically synthesized by IDT. For the multiplex SV assays, 108 gRNAs were prepared as a single pool. We used a reverse primer complementary to the tracrRNA component (same as above), and a forward primer complementary to the T7 promotor.

We have previously described our process for preparation of gRNA pools from array-synthesized oligonucleotides^14^. For this study, two ng of the input oligonucleotide pool was amplified in a 25-μl reaction mixture including 1X Kapa HiFi Hot Start Mastermix (Kapa Biosystems, Wilmington, MA) and 1-μM of each primer. The reaction was initially denatured at 95°C for 2 min, followed by 20 cycles of 20 sec of 98°C, 15 sec of 65°C and 15 sec of 72°C. The final steps for amplification involved an incubation at 72°C for 1 min and cooling to 4°C. The amplified product was purified with AMPure XP beads (Beckman Coulter, Brea, CA) in a bead solution to sample ratio of 1.8. Two hundred ng of the purified product was used as a template for *in vitro* transcription using the MEGAscript T7 transcription kit (Thermo Fisher Scientific, Waltham, MA). After the transcription reaction, the RNA products were purified using RNAClean XP beads (Beckman Coulter) in a bead solution to sample ratio of 3.0. The final gRNAs were quantified with the Qubit RNS High Sensitivity kit (Thermo Fisher Scientific). The RNA reagent kit on a LabChip GX (Perkin-Elmer, Waltham, MA) was used to confirm the product size per the manufacturer’s protocol.

### CATCH enrichment of HMW DNA molecules

The 100-kb GM12878 targets and the 200-kb *BRCA1* CATCH target were isolated using the “CATCH 100-300kb extr1h sep3h.shflow” workflow on the SageHLS instrument (Sage Science, Beverly, MA). Isolation of the 4-Mb MHC locus was performed with the “CATCH 100-300kb extr1h sep4h.shflow” workflow on the SageHLS instrument. Intact GM12878 cells (~1.5 million) were loaded into the sample well, and a lysis buffer containing 3% SDS was loaded into a reagent well just upstream of the sample well. Electrophoresis was carried out for 1 hour, thereby driving the SDS through the sample well, where the cells were rapidly lysed. With the SDS, proteins and membrane components were carried away from the sample well to the bottom electrode chamber. The genomic DNA remains very large (>>2 Megabases), and becomes firmly entangled in the agarose wall of the sample well during the extraction electrophoresis. At the end of the extraction stage, the electrophoresis was halted, the reagent well was emptied and refilled with the Cas9-gRNA reaction mixture.

For the treatment stage, a 40-μl Cas9-gRNA mixture had the following components: 1X SAGE enzyme buffer, 10-μM of the gRNA pool, and 4-μM of Cas9 enzyme (New England Biolabs, <u>Ipswich, MA</u>). The reaction was pre-incubated at 37°C for 10 min, and then mixed with 40-μl 1X SAGE enzyme buffer. Electrophoresis was carried out for one minute to drive the Cas9 enzyme into contact with the genomic DNA inside the sample well wall. Then, electrophoresis was stopped, followed by Cas9 digestion of the genomic DNA at room temperature for 30 minutes. After Cas9 digestion, the reagent well was emptied and refilled with the SDS lysis reagent, and size selection electrophoresis was carried out for three hours. The electrophoresis conditions used a pulsed field waveform designed for optimal resolution of DNA fragments 100-to 300-kb in size. After size separation, a second orthogonal set of electrodes was used to elute the size-separated DNA into a series of elution modules located along one side of the gel column. Eluted DNA was removed from the cassette within 1 hour of run termination, and the Qubit HS assay (Thermo Fisher Scientific) was used to measure the total DNA.

### Quantitation of CATCH processed DNA targets

We used a TaqMan qPCR Copy Number assays (Thermo Fisher Scientific) to measure the DNA concentration after extraction. The 10-μl reaction included 2-μl of eluted CATCH sample, 1X TaqMan Genotyping Mix, 1X TaqMan RNaseP reference, and 1X TaqMan assay for a specific target. The samples were denatured at 95°C for 10 min, followed by 50 cycles of 15 sec at 95°C and 60 sec at 60°C. For a relative quantification (i.e. target versus RNaseP reference), we used a modified ΔΔCt method^15^. 1 ng of NA12878 genomic DNA sample was used as control. For an estimation of absolute copy number in **Supplementary Figure 2**, we assumed that 290 genome copies are in 1 ng of the control sample. **Supplementary Table 10** shows the list of TaqMan Assays used for all CRISPR-linked read assays in this study.

### Library preparation, sequencing, and alignment-based phasing

Using 1.25-μl aliquot of target-enriched DNA sample from the CATCH process (typically 0.2 ng), we prepared linked read libraries using the Chromium Gel Bead and Library Kit (10X Genomics, Pleasanton, CA. We sequenced the libraries on NextSeq 500 sequencer (Illumina, San Diego, CA) with 2×151 base-pair paired-end reads using Mid Output v2 kit. With the Chromium library preparation, all resulting read pairs contain a 16-base barcode that indicates their HMW DNA molecule of origin. We used Long Ranger v2.1.6 (10X Genomics) to: i) demultiplex and convert the resulting BCL files to FASTQ files, ii) align the barcoded reads in the FASTQ files to the human genome reference build GRCh38, and iii) phase variants using the barcode information attached to each read. For Assay 2 which targets a longer region than the other assays, we optimized the parameters to get a phased block covering the entire target locus. We applied a filtering process to exclude the 1% of barcodes that had the highest read per barcode ratio, and then conducted random read downsampling (**Supplementary Table 11**).

### Local assembly and haplotype assignment

We assembled the linked reads derived from the target DNA with the Supernova assembler v2.0.0 (10X Genomics)^8^. To facilitate assembly, we removed barcodes with relatively few mapped reads or a relative abundance of mapped reads. For each barcode, the size of the genomic region spanned by the barcode was determined by the mapping locations of reads labeled with that barcode, and calculated as the difference between the largest read mapping coordinate and the smallest read mapping coordinate. For each barcode, the size of the genomic region spanned provided a proxy for the size of the original HMW DNA molecule input to the Chromium system (10X Genomics).

The size of a molecule present within a given partition can be imputed from the barcode linked sequences. For example, if one uses all the reads from a given barcode, after alignment, the general size characteristics of the originating DNA molecule are apparent. Put otherwise, the imputed size of a DNA molecule can be extrapolated from the number of mapped reads per a given barcode. However, barcodes with very few mapped reads (i.e. less than 20) do not provide enough sequences to impute these sizes. As such, barcodes with low number of reads were removed from downstream processes. Conversely, barcodes with a relatively higher abundance of mapped reads had a higher probability coming from a droplet that was occupied by both allelic copies of the target. We refer to this phenomenon as a barcode collision. To reduce the negative effect of this subset of barcodes and their associated reads on analysis, we assigned the sequence-imputed HMW molecules into 10-kb bins, i.e. 0 kb – 10 kb, 10 kb – 20 kb etc. Within each bin, we determined those barcodes with a higher probability of having a barcode collision. This was calculated by dividing the number of unique reads with the barcode by the size of genomic region spanned by the barcode. We eliminated the barcodes with the highest potential for collisions to achieve an average coverage below 70X.

After this step, we used the remaining linked reads to perform assembly. We extracted reads with these barcodes from the original FASTQ files generated by Long Ranger (10X Genomics), then used these subset FASTQ files as input to Supernova assembler (10X Genomics)^8^ to generate assembled scaffolds. The parameters used to run the assembler are indicated in **Supplementary Table 11**. To assess the scaffold structures of the Supernova output, we compared where each raw FASTQ read aligned to the reference genome GRCh38 with where it aligned to the assembled scaffold; plotting this comparison in R provided a visual representation of scaffold structure (**Fig. 2a, Supplementary Fig. 8**).

### Haplotype validation

To assess the quality of Supernova assemblies for each haplotype, we compared the allelic content of the assembled scaffolds with a data set considered as the ground truth haplotypes. To determine the allelic content of our assembled scaffolds, we aligned each scaffold to the human reference genome GRCh38 using minimap2 v2.7 and then generated a VCF file of variant calls using samtools mpileup v1.6^16^. The ground truth data set for comparison was the Platinum Genome phased variants (downloaded from: ‘ftp://ussd-ftp.illumina.com/2017-1.0/hg38/hybrid/hg38.hybrid.vcf.gz’). The ground truth variant dataset was filtered to obtain only phased heterozygous SNVs. We then compared the shared positions between the CRISPR-linked read scaffold variant calls and the filtered ground truth data sets, and calculated: number of shared alleles between the datasets / number of shared variant positions between the datasets. We expected that each assembled scaffold should share all of its alleles with one of the two ground truth haplotypes in the target genomic region.

Additionally, this process was also used to assign haplotypes to assembled scaffolds when the assembly process failed to generate a single contiguous scaffold for a target locus (**Supplementary Figure 2**). In this case, instead of the ground truth haplotypes, the haplotypes obtained by the alignment-based process (i.e. the output of Long Ranger) was used for the comparison.

### Comparison with other assemblies

We compared our assembled scaffolds with assemblies generated from long read sequencing and linked read whole genome sequencing. The long-read assemblies were a Pacific Biosciences assembly^1^ (downloaded from: ‘ftp://ftp.ncbi.nlm.nih.gov/genomes/all/GCA/001/013/985/GCA_001013985.1_ASM101398v1/GCA_001013985.1_ASM101398v1_genomic.fna.gz’) and an Oxford Nanopore assembly^2^ (downloaded from: http://s3.amazonaws.com/nanopore-human-wgs/canu.30x.contigs.fasta). Both of these assemblies were haploid and thus included only one sequence for each genomic region. To locate where the sequences from our targeted assays occurred in these other assemblies, we aligned GRCh38 target sequences to the assemblies using minimap2 v2.7^17^. Alignment of our target sequences provided a unique location for each target (**Supplementary Table 1** for Assays 1 and 2, **Supplementary Table 6** for Assay 3). To obtain the structure and allelic content of the assemblies, we used the same method as described in ‘Local assembly and haplotype assignment’ and ‘Haplotype validation’.

### Identification of HLA genotypes

All the reported HLA gene alleles (downloaded from: ‘ftp://ftp.ebi.ac.uk/pub/databases/ipd/imgt/hla/hla_gen.fasta.txt’) were aligned to our assembled scaffolds using minimap2 v2.7^17^. Among the alignments having a percent match greater than 90%, we selected the closest allele sequence considering both the edit distance and overall length of insertions and deletions, which were provided in the alignment output.

## REFERENCES

1. Pendleton, M. et al. Assembly and diploid architecture of an individual human genome via single-molecule technologies. Nature methods 12, 780–786 (2015).

2. Jain, M. et al. Nanopore sequencing and assembly of a human genome with ultra-long reads. Nature biotechnology 36, 338–345 (2018).

3. Bell, J.M. et al. Chromosome-scale mega-haplotypes enable digital karyotyping of cancer aneuploidy. Nucleic acids research 45, e162 (2017).

4. Greer, S.U. et al. Linked read sequencing resolves complex genomic rearrangements in gastric cancer metastases. Genome medicine 9, 57 (2017).

5. Zheng, G.X. et al. Haplotyping germline and cancer genomes with high-throughput linked-read sequencing. Nature biotechnology 34, 303–311 (2016).

6. Xia, L.C. et al. Identification of large rearrangements in cancer genomes with barcode linked reads. Nucleic acids research 46, e19 (2018).

7. Jiang, W. et al. Cas9-Assisted Targeting of CHromosome segments CATCH enables one-step targeted cloning of large gene clusters. Nature communications 6, 8101 (2015).

8. Weisenfeld, N.I., Kumar, V., Shah, P., Church, D.M. & Jaffe, D.B. Direct determination of diploid genome sequences. Genome Res 27, 757–767 (2017).

9. Eberle, M.A. et al. A reference data set of 5.4 million phased human variants validated by genetic inheritance from sequencing a three-generation 17-member pedigree. Genome research 27, 157–164 (2017).

10. Complete sequence and gene map of a human major histocompatibility complex. The MHC sequencing consortium. Nature 401, 921–923 (1999).

11. Moore, J.K. & Haber, J.E. Cell cycle and genetic requirements of two pathways of nonhomologous end-joining repair of double-strand breaks in Saccharomyces cerevisiae. Molecular and cellular biology 16, 2164–2173 (1996).

12. McVey, M. & Lee, S.E. MMEJ repair of double-strand breaks (director’s cut): deleted sequences and alternative endings. Trends in genetics : TIG 24, 529–538 (2008).

13. Schneider, V.A. et al. Evaluation of GRCh38 and de novo haploid genome assemblies demonstrates the enduring quality of the reference assembly. Genome research 27, 849–864 (2017).

14. Shin, G. et al. CRISPR-Cas9-targeted fragmentation and selective sequencing enable massively parallel microsatellite analysis. Nature communications 8, 14291 (2017).

15. Livak, K.J. & Schmittgen, T.D. Analysis of relative gene expression data using real-time quantitative PCR and the 2(-Delta Delta C(T)) Method. Methods 25, 402–408 (2001).

16. Li, H. et al. The Sequence Alignment/Map format and SAMtools. Bioinformatics 25, 2078–2079 (2009).

17. Li, H. Minimap2: pairwise alignment for nucleotide sequences. Bioinformatics (2018).

